# Genome-wide analysis reveals mutual gene flow between goats from Island Southeast Asia and from Southern Africa

**DOI:** 10.1101/2025.06.13.659639

**Authors:** Ryo Masuko, Ayin, Maho Masaoka, Fuki Kawaguchi, Shinji Sasazaki, Muhammad I. A. Dagong, Sri R. A. Bugiwati, Joseph S. Masangkay, Jiaqi Wu, Takahiro Yonezawa, Johannes A. Lenstra, Hideyuki Mannen

**Affiliations:** Laboratory of Animal Breeding and Genetics, Graduates School of Agricultural Science, Kobe University, Kobe, Japan; Faculty of Animal Science, Hasanuddin University, Makassar 90245, South Sulawesi, Indonesia; College of Veterinary Medicine, University of the Philippines, Los Baños, Philippines; Graduate School of Integrated Sciences for Life, Hiroshima University, Japan; Department of Molecular Life Science, Tokai University School of Medicine, Japan; Faculty of Veterinary Medicine, Utrecht University, Netherlands

## Abstract

This study aimed to reveal a comprehensive genetic diversity and genetic structure of Island Southeast Asian (ISEA) goats, and infer the details of gene introgressions and propagation routes in ISEA goats using genome-wide SNPs. Correlation analysis of the distance from the domestication center and the genetic diversity of each Asian population showed a significant negative correlation (r = - 0.796, p = 2.66E-05), while the Philippines showed a relatively high genetic diversity (He = 0.363) against the distance, suggesting the multiple propagation routes and admixture history in Philippine goats. Reduced Representation Admixture Analysis at K=6 showed that six clusters (Europe, Northwest Africa, Southern Africa, Eastern Africa, Asia and Southeast Asia) were formed based primarily on geographical location, and ISEA goats included African and Europe components. However, this introgression is much lower in Indochinese goats. In addition, the Southern African and European populations included Southeast Asia component. Treemix and f4 statistics analyses showed the genetic influence and gene flow between ISEA and the Southern Africa. The results of this study have revealed obvious mutual gene flow between ISEA and the Southern African regions through the European maritime activities across the Indian Ocean since the Middle Ages.

## Introduction

Since their domesticated in the Fertile Crescent approximately 11,000 years ago before present, goats (*Capra hircus*) have become widespread throughout the world [1]. Because of their small size and adaptability to the environment [2,3], they are one of the livestock species that could be easily transported across land and maritime following human movement. The genetic information of goats is important for elucidating genetic relationships, origins of domestication and history of propagation, and has been studied using mtDNA, SRY gene on the Y chromosome, and single nucleotide polymorphisms (SNPs) [4–9].

Previous studies using mtDNA control region sequences revealed six maternal haplogroups (A, B, C, D, F, and G) [9–13]. The haplogroup A is the most widespread in worldwide, while minor haplogroups C, D, F, and G are sporadically found at low frequencies in Asia, Europe, the Near East, and North Africa, respectively [12–15]. The mtDNA haplogroups have been reported to show a weak phylogenetic structure due to extensive intercontinental transportation of goats in the past [12]. Whereas, the haplogroup B is found at high frequency in Southeast Asia. Given the unique phylogeographic distribution of the haplogroup B, the domestication of goats is thought to have also occurred in further east of the Fertile Crescent [11,16], but the origin and propagation route are still unclear. In contrast to the mtDNA haplogroups, the Y-chromosomal haplotypes (Y1AA, Y1AB, Y1B, Y2A, and Y2B) show a clear difference in geographic structure between continents due to bottlenecks and expansions caused by the goat migration process [17]. Y1AB dominating in northern China and Y1AA with Y2B in the south. Y1B, Y2A, and Y2B have been observed mainly in Europe, Africa, and East Asia, respectively [9,17–19].

Since the development of genome wide SNP Chip for goats [20], studies have been conducted to assess genetic diversity and to estimate propagation routes for goats worldwide [6,8,21–26]. Autosomal 55K SNP [6] and whole genome sequence [27] studies show a strong regional difference of the genetic structure similar to SRY haplotypes. In Southeast Asian goats, previous studies reveal that similar genetic diversity between Lao and Chinese/Mongolian goats [24], and multiple propagation routes from domestication center to the Mainland Southeast Asia (MSEA) [26].

The Philippines and Indonesia, located in Southeast Asia, are archipelagic nations consisting of several islands. These Island Southeast Asia (ISEA) are known as mega-diversity countries with a wide variety of species. In the ISEA, there are records of trade and Islamic missionary work by Muslim merchants from the 9th century and colonial rule by Spain in the Philippines and the Netherlands in Indonesia from the 16th century [28–30]. In maternal lineage analysis of ISEA goats, the low frequency of haplogroup A is demonstrated in Indonesia [31,32] and two propagation routes to ISEA through the Asian continent are estimated by inference on population dynamics [32]. It is also inferred gene introgression through the Indian ocean between ISEA and Europe or Africa from the comprehensive result of maternal and paternal lineage analyses: the distribution of mtDNA haplogroup B in the southern African region and the presence of the SRY haplotype Y2A in ISEA which is predominant in Europe and Africa but absent in MSEA [32].

These gene introgressions across the Indian Ocean are thought to be associated with human trade activities since the Middle Ages. Therefore, detailed information in ISEA goats by genome-wide survey is expected to reveal ISEA specific gene structure and past human trade activities. Hence, this study focused on the ISEA goats with seven islands and six regions, and aimed to (1) reveal a comprehensive genetic diversity and genetic structure of ISEA goats, and (2) infer the details of gene introgressions and propagation routes in ISEA goats through the Indian Ocean using genome-wide SNP information on ISEA goats.

## Materials and Methods

### Ethics declarations

This study is reported in accordance with ARRIVE guidelines (https://arriv eguid elines.org). All experiments were carried out according to the Kobe University Animal Experimentation Regulations, and all protocols were approved by the Institutional Animal Care and Use Committee of Kobe University and by Association for the Promotion of Research Integrity (Approval Number: AP0000436777). All blood samples collections were approved by animal owners with signed informed consent.

### Sample Collection and Genotyping

We obtained DNA samples 168 native goats from 8 countries: Philippines (n = 57), Indonesia (n = 30), Vietnam (n = 7), Laos (n = 5), Cambodia (m = 25), Myanmar (n =5), Bangladesh (n = 34), Bhutan (n =5) (Fig 1a, S1 Table). According to previous study [26], Cambodian goats were divided into Cambodia-M in a remote mountainous area east of the Mekong and Cambodia-P in the plain west of the Mekong, which clearly differ in morphology [26] and in the frequency of the mtDNA haplogroup B [11]. Similarly, Myanmar goats were divided into the Myanmar and Myanmar-Shan populations [26]. All goats were genotyped using Illumina goat SNP 50K Bead chip (53,347 SNPs) (Illumina, Inc. San Diego, CA, USA).

**Fig 1.**
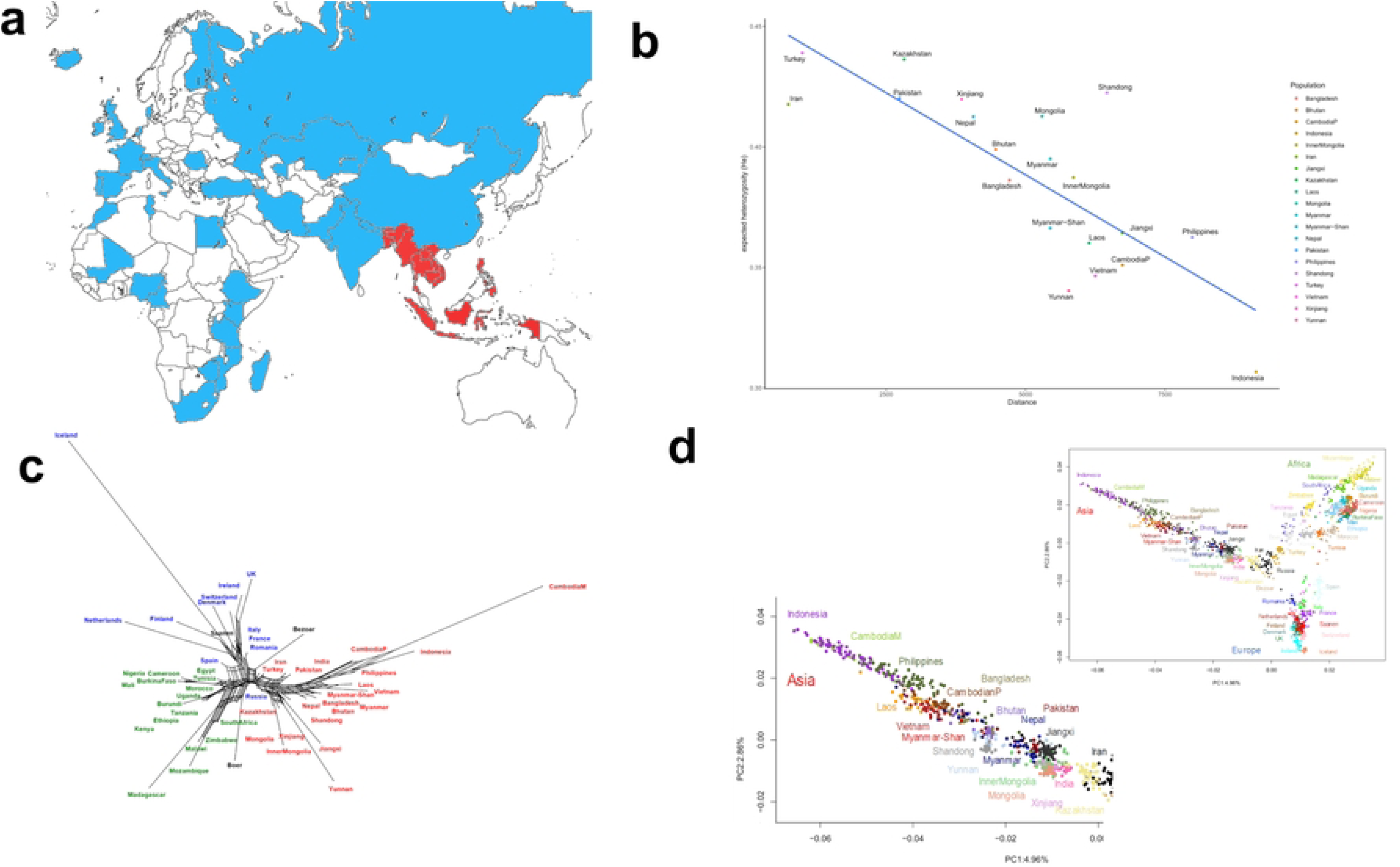
Genetic diversity in Asian populations and the relationship among the Old-World goats. (a)The Old-World goats used in this study. Red and blue color indicate locations of goat populations used in this study and previous studies [6,21,22,26,33], respectively. The map was generated using Inkscape 1.4 (86a8ad7, 2024-10-11) based on a free map website by 3kaku-K (https://www.freemap.jp/item/world/world1.html). (b)The correlation between expected heterozygosity of Asian goat population removing CambodiaM and distance between their sampling locations and the domestication centers. This plot was generated using RStudio 2023.09.0+463 “Desert Sunflower”. (c) Neighbor-Network of the Old-World goats using SplitsTree App. The edge lengths are proportional to the pairwise FST distances between populations: Blue, green, and red colors showed Europe, Africa, and Asian populations, respectively. (d) Supervised PCA of the Old-World goats and the enlarged plot focused the Asian goats. These plots were generated by using RStudio 2023.09.0+463 “Desert Sunflower”.

### Dataset Construction and quality control

In this study, two datasets (Dataset1 and Dataset2) were constructed for use in calculating genetic diversity and other analyses, respectively. We combined our data with the previously published goat 50K SNP data sets [6,21,22,26,33], but without the cosmopolitan Angora, commercial breeds in South Africa (Kalahari Reds and Savanna), the New World, and the Oceanian populations. The resulted in genotypes of 4,480 goats from 45 countries and seven wild bezoars from Iran for 44,002 SNPs. After merging, the quality control was performed using plink v1.9. We excluded SNPs with call rate < 0.02, maf < 0.001, removing individuals with call rates < 0.02 as well as individuals identified as outliers and LD pruning (plink indep-parwise: window size 50Kb, step size 5, r^2^ threshold 0.04). This procedure left 21,009 SNPs with 4,318 individuals representing 54 wild or domestic populations from 45 countries for the analysis. We defined this dataset as “Dataset1”. For “Dataset2”, population with more than 40 goats except ISEA population were randomly sampled to 40 individuals per population to reduce ascertainment bias. The “Dataset2” consisted of 21,009 SNPs with 1,747 goats/bezoar (S1 Table).

### Genetic diversity and population structure analysis

Observed and expected heterozygosity (Ho and He) for each SNP were calculated using plink 1.9 [34]. The most probable domestication centers for goats are Ganj Dareh (34°16’19.56 N, 47°28’32.88 E) in West Zagros of Iran and Nevali Cori (37°31’06.0”N 38°36’20.0”E) in Taurus of Turkey [12,35–37]. Therefore, we calculated the correlation between the He and the average geographic distances (in kilometers) to the domestication centers from the capital of each population. Principal Component Analysis (PCA) and supervised Principal Component Analysis (svPCA) was done with plink 1.9 and plotted by R 4.2.3 [38].Genetic population structure and admixture using dataset2 were estimated using ADMIXTURE version 1.3.0 [39] with K=1 to 15, 20, 30, and 40. The RRAA (reduced representation Admixture analysis) was performed to suppress the inbreeding bias by reducing the representation of inbred breeds [26]. We implemented by random selecting six individuals from Cambodia M population, running the Admixture for 100 times of such selections and averaging for each individual the percentages of the inferred genomic structure.

### Phylogeographic inference and admixture events

To assess the differentiation within and across breeds, pairwise weighted F_ST_ distances between breeds were calculated with R package StAMPP and visualized in Neighbor-Net graphs using SPLITSTREE [40]. Admixture among populations was inferred by the TREEMIX 1.13 [41] with m=0 to 15 migration edges, and bezoars as outgroup. In Treemix analysis, we defined clusters goat populations into regions based on geographical locations and the RRAA pattern: Europe, Northwest Africa, Southern Africa, Eastern Africa, Boer, West Asia, Central Asia, South Asia, East Asia, MSEA, Philippines, Indonesia (S1 Table). To estimate shared evolutionary history between ISEA goats and other populations, we also performed outgroup f3 statistics and f4 statistics using R package ADMIXTOOLS 2 ver 2.04 [42,43]. These statistics were computed in the forms of f3(O; X, Y) and f4(O, X; Y, Z), where O represents the outgroup (Bezoar), Y is the ISEA populations, and X and Z are other populations. For f4 statistics, statistical significance was determined at a threshold of p < 0.001.

## Results

### Genetic diversity and phylogenetic analysis

The regional Ho and He were calculated for the ISEA (S2 Table), indicating that the values for Philippines goats (Ho: 0.328; He: 0.363) are higher than those of the Indonesian (Ho: 0.278; He: 0.307). For seven regions in ISEA, the Ho ranged from 0.278 (Sulawesi) to 0.344 (Samar & Leyte) and He ranged from 0.307 (Sulawesi) to 0.355 (Samar & Leyte). In Asian populations, Cambodia-M showed quite low genetic diversity (He: 0.158). We also investigated the correlation between the He of Asian populations and the average distance from the sampling locations to the domestication centers. The Correlation coefficient of Asian populations showed significant (r = −0.593, p = 0.00461) (S1 Fig). In case that the value of Cambodia-M is removed as an outlier, highly significant correlation (r = −0.796, p = 2.66E-05) was obtained (Fig 1b). The Philippine and Shandong populations showed relatively high diversity given their long geographic distance from the domestication center.

To infer the genetic relationship among Old World populations, pairwise F_ST_ distance were calculated (S3 Table). The genetic distance for the ISEA goats showed the lowest value between Philippines and Indonesia (0.0716) and the greatest value between Iceland and ISEA (Philippines: 0.381; Indonesia: 0.456), and generally reflected a correlation with geographic distance. The Neighbor-Network based on pairwise F_ST_ distance showed the correlation with geographic region and clearly separated Asia, Africa and Europe by the network bottleneck (Fig 1c). ISEA populations formed a distinct clade together with Cambodian populations from MSEA populations. The long edges were observed in the Iceland, Madagascar, and Cambodia-M populations, indicating high inbreeding by geographic isolation like recent studies [6,26].

### Population and Genetic Structure analysis

The svPCA was performed to avoid the influence of high inbreeding in Cambodia-M as described in our previous study [26]. The Cambodia-M located as forefront independent population in the unsupervised PCA (S2 Fig) but clustered with Southeast Asian (SEA) populations in the svPCA (Fig 1d). The svPCA identified three distinct clusters, Asia, Africa, and Europe centering on the Middle East populations in domestication center. PC1 (4.96%) distinguished Asia from Europe and Africa, while PC2 (2.86%) separated Europe from Asia and Africa, respectively. ISEA goats clustered with MSEA goats, and most of Indonesian goats were genetically similar to Cambodia-M, whereas the Philippines goats resemble those from Laos and Vietnam despite being separated by an ocean.

The ADMIXTURE plots showed Asian, African, and European clusters at K=3 (S3 Fig), similar to the results of PCAs and phylogenetic analysis. At K≥4, ISEA populations were separated from SEA and Cambodia-M population showed independent genetic structure. Although it looks like the ancestral component, it is also possible the influence of inbreeding bias by Cambodia-M causes this genetic structure. To reduce this bias, we performed RRAA at K=2 to 8 (S4 Fig). RRAA plots are similar to the ADMIXTURE plots at K=2 to 4, whereas a reduction of the inbreeding bias by Cambodia-M shows that the ISEA populations cluster with other SEA populations at K=5 to 8 (Fig 2a).

**Fig 2.**
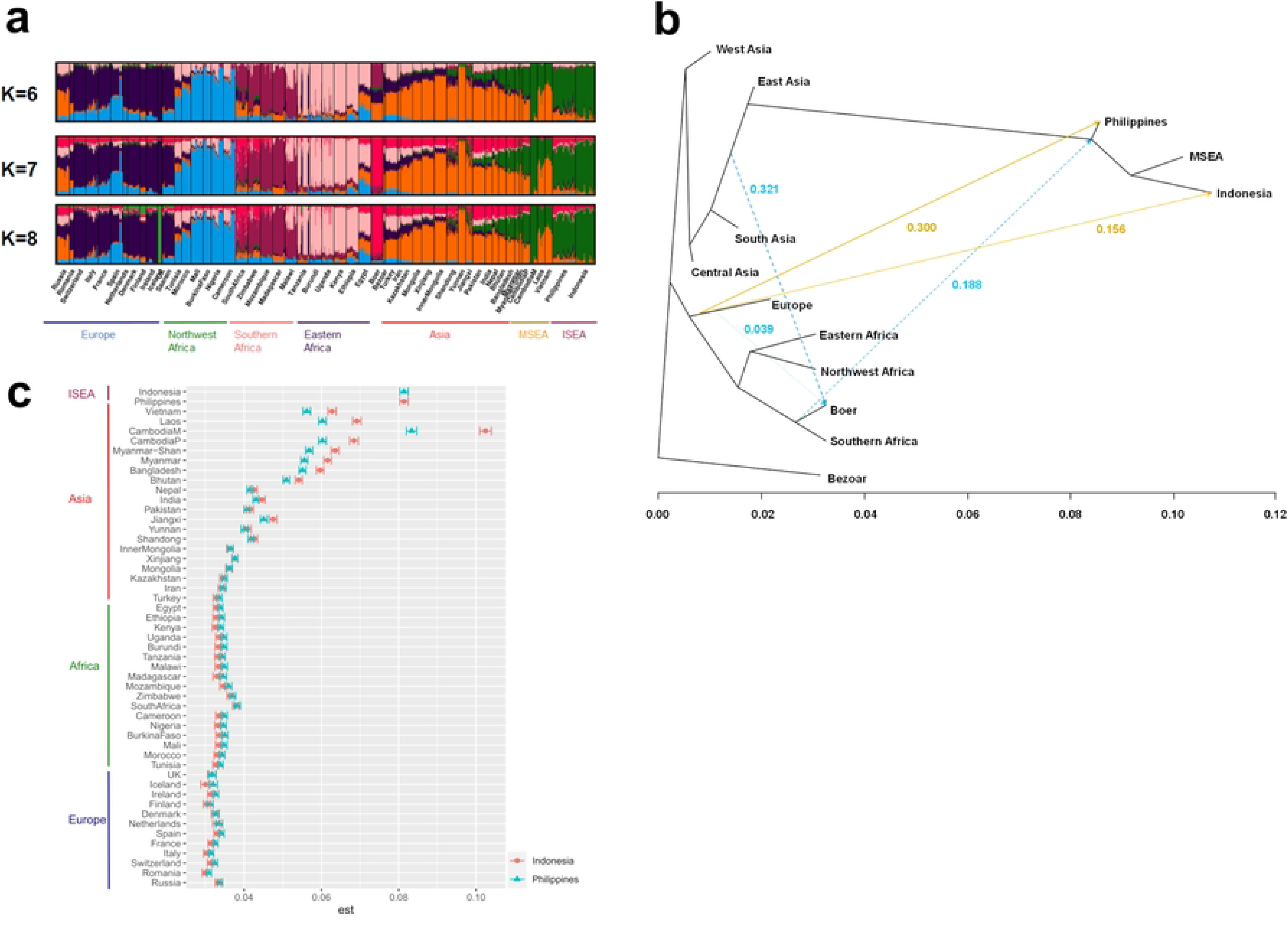
Population genomic structure and gene flow among the Old-World goats. a) Reduced Representation Admixture analysis (RRAA) by the ADMIXTURE program ver. 1.3 (K=6-8). This figure was generated using CLUMPAK (https://clumpak.tau.ac.il/). Abbreviations: SEA, Southeast Asia; MSEA, Mainland Southeast Asia; ISEA, Island Southeast Asia. b) Regional Treemix graphs at m=13 of the Old-World goats. Only edges related to ISEA (orange line) and Boer (sky blue broken line) are shown with migration weights. c) The f3-statistics between ISEA goats and other domesticated goats. This plot was generated using RStudio 2023.09.0+463 “Desert Sunflower“.

The RRAA (K=6 to 8) showed that European populations exhibited a homogeneous genetic structure, while African populations were divided into three genetic structures based on the geographic regions: Northwest Africa (light blue), Southern Africa (dark violet), and Eastern Africa (pink) (Fig 2a). Asian populations showed the geographical cline by Asian component (orange) and SEA component (green). Based on the RRAA at K=6, components of Northwest Africa (light blue), Southern Africa region (dark violet), Eastern Africa (pink) and Europe (dark blue) were observed in ISEA goats, especially the Philippine goats at the relatively high proportions (1.76% - 7.08%. The total proportion of European and African components is 17.03%) but is much lower in Indochinese goats (Cambodia-M: 0.10%, Laos: 2.36%, and Vietnam: 1.17%) (S4 Table). Notably, SEA component was observed in the Southern African and European goats, especially in South Africa (6.10%) and Zimbabwe (4.37%) compared with another African regions (S4 Table). RRAA at K=7 and 8 showed a Boer component (deep pink) which were observed in most of Asian countries including ISEA (Fig 2a and S5 Table), but rarely in Indochinese goats (Cambodia-M: 0.0%, Laos: 1.1%, and Vietnam: 0.1%).

### Migration analysis

Treemix analysis was performed to investigate gene flow from/to ISEA and Boer. At m=13 (Fig 2b, S5 Fig), Treemix graph clustered the Old-World goats according to their geographic origin. Migration edges were observed from the root of European/African populations to the tip of Philippines (migration weight: 0.300) and Indonesia (0.156), and from near the root of Boer/Southern Africa and SEA (0.188). In addition, Boer goats showed migration edges between Asia, Africa and Europe, which is consistent with establish process of the Boer [44]. It should be noted that Treemix may occasionally infer an incorrect direction of migration [45].

In order to infer the admixture and migration and to confirm the confidence of the Treemix result, outgroup f3 and f4 statistics were calculated for pairs of ISEA and other domesticated goat populations. Outgroup f3 statistics (Bezoar; ISEA goats, other populations) are generally proportional to geographic distance with ISEA (Fig 2c). Interestingly, South Africa has relatively high values (Philippines: 0.03830; Indonesia: 0.03782) and showed a close genetic distance with ISEA goats. We estimated f4 (Bezoar, Africa/Europe population; ISEA goats other domestication goats) and selected the pairs with significant value (p-value<1.00e-04) because Treemix suggested migration between ISEA and South Africa/Europe region (S6 Table). ISEA goats shares ancestry with Europe and Africa (Philippines: Romania, Switzerland, Italy, France, Spain, Netherlands, Denmark, Finland, Ireland, Iceland, UK, Tunisia, Morocco, Mali, Burkina Faso, Nigeria, Cameroon, South Africa, Zimbabwe, Mozambique, Malawi, Tanzania, Burundi, Uganda, Kenya, Ethiopia; Indonesia: Switzerland, Italy, France, Spain, Netherlands, Denmark, Finland, Ireland, UK, Burkina Faso, Nigeria, Cameroon, South Africa, Zimbabwe). In European and African goat populations, South Africa showed a stronger genetic association with ISEA than with European goats (Russia, Romania, Switzerland, Italy, France, the Netherlands, Denmark, Finland, Ireland, UK). (S6 Table).

## Discussion

The ISEA goats had low genetic diversity compared with the heterozygosity of MSEA (S2 Table), suggesting the influence of a bottleneck due to geographical isolation as islands and the reflection of geographical distance from the domestication center. This lower genetic diversity of ISEA goats is consistent with the results of our previous study using mtDNA [32]. The PCAs and Neighbor-Net illustrated consistently differentiate goats based on the geographical distance from the domestication center (Figs 1c and 1d, S2 Fig). The svPCA also demonstrated that Philippine goats clustered MSEA goats, whereas most of the Indonesian goats are genetically slightly more related to the Cambodia-M population. In contrast to the previous report in Asia [46], the He values in Asia showed strong significant correlation with the distance from the domestication centers (Fig 1b), which is probably due to the sufficient number of various populations in this study. While the He of the Philippines showed a relatively high value against the distance in ISEA. These results of the svPCA and the He value in ISEA were probably reflected the admixture status of mtDNA haplogroups A and B in the Philippines [32]. In other words, the haplogroup A arrived in the Philippines via China/Taiwan, whereas goats with the haplogroup B were introduced from the Malay Peninsula to the Philippines via Indonesian islands and subsequently genetic admixture events occurred in the Philippines [32].

Intriguingly, RRAA, Treemix, and f4 statistics analyses showed the genetic influence and gene flow between ISEA and Europe/Africa (Figs 2a and 2b, S6 Table). Treemix showed migration edges between ISEA and Southern Africa/Europe, suggesting gene flow events among these regions (Fig 2b). Notably, the f4 statistics revealed that European and African populations shared a minor ancestry with ISEA. In addition, South Africa population had a relatively stronger genetic relationship with ISEA than with European populations (S6 Table). RRAA analysis at K = 6 to 8 showed higher proportions of European and African components in both Indonesian and, slightly more, Philippine goats (Fig 2a, S4 Table). Focusing on the details of the RRAA at K = 6, the SEA component (green) was shown in the Southern African populations, especially in South Africa (6.10%). This genetic component was lower in other African populations (<2.0%) (Fig 2a, S4 Table), suggesting the gene introgression from SEA to the Southern Africa occurred via the Indian Ocean. Based on the results of Treemix and f4 statistics, the SEA component shown in the Southern Africa was probably due to the genetic influence of gene flow from ISEA goats.

The propagation of domesticated animals between geographically distant regions can be linked to human migrations [47–49]. During the Age of Discovery from the 15th century, European countries conducted maritime trade in the Indian Ocean, and there are also historical records of chickens, sheep, and goats being transported aboard ships as a source of protein during voyages [50,51]. Since goat’s milk was used as a protein source during voyages, both female and male goats seem to have been transported aboard ships. The coastal regions of Africa were under European colonial rule and were used as relay points in maritime trade. During this period, the Philippines was colonized by the Spanish and used as an important hub for trade with Asia [30]. In Indonesia, the spice trade was actively conducted by European countries under the colonial rule of the Dutch or Portuguese [52,53]. These historical backgrounds were probably reflected in the genetic relationships and gene flow between ISEA and Southern Africa, as suggested in this study (Figs 2a and 2b, S6 Table).

In addition, the South Africa which was suggested in this study to have a relatively strong association with ISEA, had historical backgrounds related to maritime trade by European countries. In South Africa, the Cape Colony was established by the Dutch East India Company in the 17th century and used as a port of call for Oriental trade [54]. Subsequently, the British colonized South Africa from the 18th century. These historical backgrounds suggest that the genetic relationship between ISEA and Southern Africa, particularly South Africa, demonstrated in this study, is a mutual gene flow through maritime trade in the Indian Ocean by European countries. This also explain the South-African – Asian crossbred origin of the Boer goat, which emerged in South-Africa during the 19^th^ century and was exported in the 20^th^ century to several Asian countries (Fig 2a) [44].

In previous maternal lineage analysis, a distinct SEA haplogroup B was observed in Tanzania, South Africa and Namibia [13,55]. The haplogroup B is also absent further north than Tanzania in Africa, suggesting gene flow from SEA region to Southern/Eastern Africa via the Indian Ocean. Based on the mutual gene introgression between ISEA and the Southern African regions suggested in this study using genome-wide analysis, the distribution of mtDNA haplogroup B in the Southern/Eastern African region probably also reflected the gene introgression from the ISEA related goats brought back from ISEA by maritime transport.

RRAA at K = 7 and 8 revealed a clear genetic component of Boer goats in most Asian breeds but not in Indochina and Madagascar (Fig 2b, S5 Table). This may reflect the import of global breeds, such as dual purpose AngloNubian and Boer meat goats in the world [44]. However, Treemix showed no direct migration edge between ISEA and Boer (Fig 2b), and the Y-chromosomal haplogroup Y2A in ISEA goats has a high frequency in Africa but not in the Boer goats with a Y1AA haplogroup [17,32]. The recent hybrid (Africa × Asian) origin of Boer complicates the analysis of its introgression in other populations and the Boer component in ISEA inferred by RRAA analysis may also represent the introduction of the other European, Asian and African goats. Further research is needed on the global genetic impact of the Boer goats.

In this study, the different genetic structure and the degree of the gene introgression was elucidated among Philippine and Indonesian goats, suggested the different propagation routes within ISEA. In addition, the high genetic diversity of Philippine goats in ISEA strengthened our hypothesis that the two propagation routes from MSEA and subsequently genetic admixture occurred in the Philippines. Considering that the transport of domesticated animals between geographically distant regions can be linked to human migrations, we suggest a mutual gene flow between ISEA and Southern African region through European maritime transport across the Indian Ocean (Fig 3). The genetic information obtained in this study will provide important insights into genetic resources of Southeast Asian goats.

**Fig 3.**
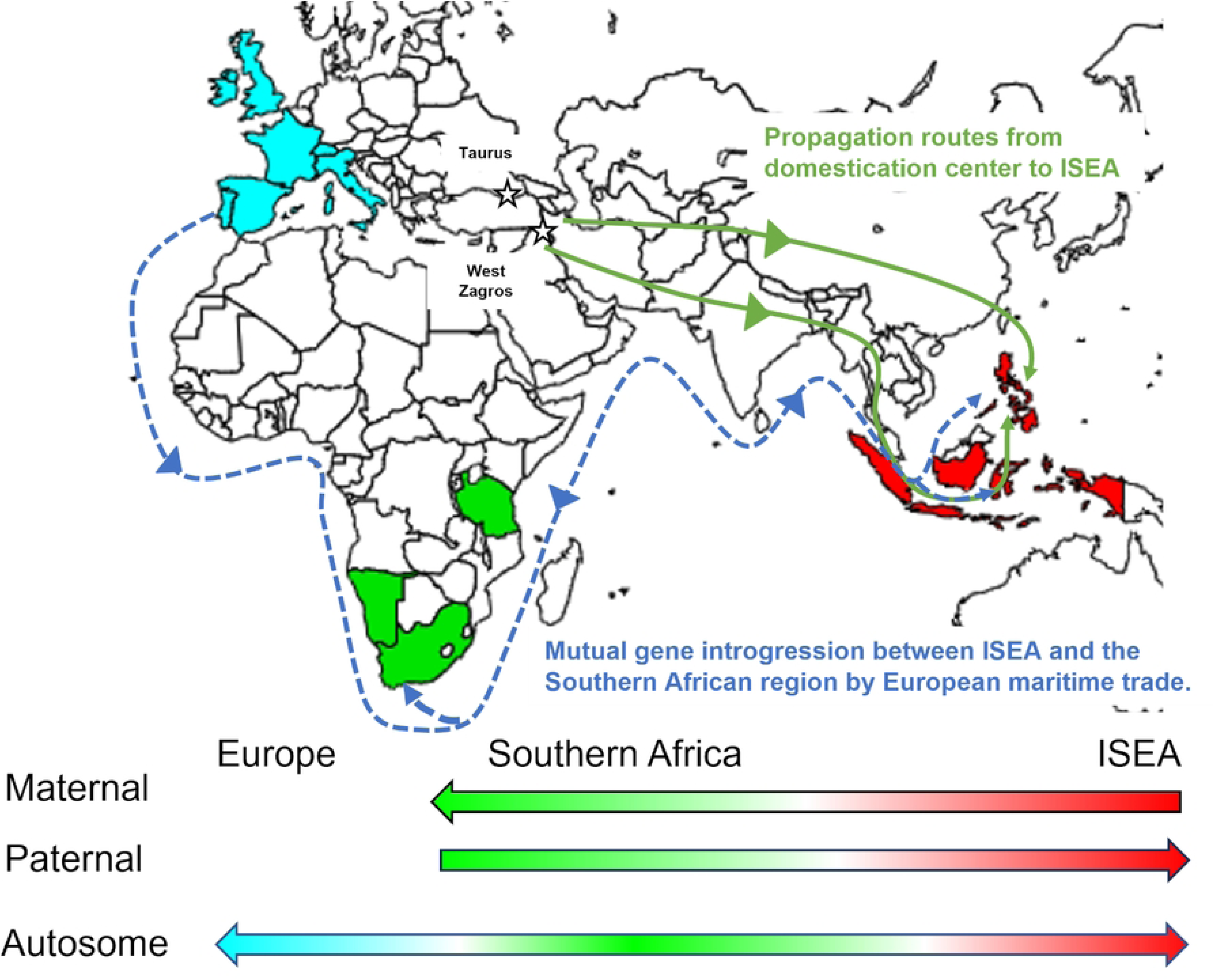
The gene introgression through the Indian Ocean. Red, green and blue colors show ISEA, the Southern African, European region, respectively.

Green solid line shows the propagation routes from the domestication center to ISEA [32]. Blue dot line shows the migration route based on historical voyage recode and mutual gene introgression between ISEA and the Southern African region. The map was generated using Inkscape 1.4 (86a8ad7, 2024-10-11) based on a free map website by 3kaku-K (https://www.freemap.jp/item/world/world1.html).

## Acknowledgments

N/A

## Supporting information

**S1 Table. Sample List of Datset1 and Dataset 2. ○ shows samples used in Dataset 2.**

**S2 Table. Expected (He) and observed (Ho) heterozygosity in Asian goats.**

**S3 Table. Pairwise FST distances among the Old World goat populations.**

**S4 Table. Proportion of the genetic components in each population in Reduced Representation Admixture analysis at K=6.**

**S5 Table. Proportion of the Boer component in each population in Reduced Representation Admixture analysis at K=7 and 8.**

**S1 Fig. The correlation between expected heterozygosity of Asian goat populations and distance between their sampling locations and the domestication centers.** This plot was generated using RStudio 2023.09.0+463 “Desert Sunflower”.

**S2 Fig. Unsupervised principal component analysis (PCA) on the Europe, Africa, Asian domestic goats and bezoars.** This plot was generated using RStudio 2023.09.0+463 “Desert Sunflower“.

**S3 Fig. Normal Admixture analysis of Eurasian and African goats at K=2 to 15, 20, 30, and 40 by the ADMIXTURE program ver. 1.3.** This figure was generated using CLUMPAK (https://clumpak.tau.ac.il/).

**S4 Fig. Reduced Representation Admixture analysis (RRAA) of the Eurasian and African goats by the ADMIXTURE program ver. 1.3 (K=2-8).** This figure was generated using CLUMPAK (https://clumpak.tau.ac.il/).

**S5 Fig. Treemix analysis of Eurasian and African goats at m=13.**

## Reference

1. Daly KG, Maisano Delser P, Mullin VE, Scheu A, Mattiangeli V, Teasdale MD, et al. Ancient goat genomes reveal mosaic domestication in the Fertile Crescent. Science. 2018;361: 85–88. doi:10.1126/science.aas9411

2. Pereira F, Amorim A. Origin and Spread of Goat Pastoralism. 1st ed. Encyclopedia of Life Sciences. 1st ed. Wiley; 2010. doi:10.1002/9780470015902.a0022864

3. Zheng Z, Wang X, Li M, Li Y, Yang Z, Wang X, et al. The origin of domestication genes in goats. Science Advances. 2020;6: eaaz5216. doi:10.1126/sciadv.aaz5216

4. Cinar Kul B, Bilgen N, Lenstra JA, Korkmaz Agaoglu O, Akyuz B, Ertugrul O. Y-chromosomal variation of local goat breeds of Turkey close to the domestication centre. J Anim Breed Genet. 2015;132: 449–453. doi:10.1111/jbg.12154

5. Colli L, Lancioni H, Cardinali I, Olivieri A, Capodiferro MR, Pellecchia M, et al. Whole mitochondrial genomes unveil the impact of domestication on goat matrilineal variability. BMC Genomics. 2015;16: 1115. doi:10.1186/s12864-015-2342-2

6. Colli L, Milanesi M, Talenti A, Bertolini F, Chen M, Crisà A, et al. Genome-wide SNP profiling of worldwide goat populations reveals strong partitioning of diversity and highlights post-domestication migration routes. Genet Sel Evol. 2018;50: 58. doi:10.1186/s12711-018-0422-x

7. Mannen H, Nagata Y, Tsuji S. Mitochondrial DNA Reveal that Domestic Goat (Capra hircus) are Genetically Affected by Two Subspecies of Bezoar (Capra aegagurus). Biochemical Genetics. 2001.

8. Somenzi E, Senczuk G, Ciampolini R, Cortellari M, Vajana E, Tosser-Klopp G, et al. The SNP-Based Profiling of Montecristo Feral Goat Populations Reveals a History of Isolation, Bottlenecks, and the Effects of Management. Genes. 2022;13: 213. doi:10.3390/genes13020213

9. Waki A, Sasazaki S, Kobayashi E, Mannen H. Paternal phylogeography and genetic diversity of East Asian goats. Animal Genetics. 2015;46: 337–339. doi:10.1111/age.12293

10. Joshi MB, Rout PK, Mandal AK, Tyler-Smith C, Singh L, Thangaraj K. Phylogeography and Origin of Indian Domestic Goats. Molecular Biology and Evolution. 2004;21: 454–462. doi:10.1093/molbev/msh038

11. Lin BZ, Kato T, Kaneda M, Matsumoto H, Sasazaki S, Mannen H. Genetic diversity and structure in Asian native goat analyzed by newly developed SNP markers. Animal Science Journal. 2013;84: 579–584. doi:10.1111/asj.12039

12. Luikart G, Gielly L, Excoffier L, Vigne J-D, Bouvet J, Taberlet P. Multiple maternal origins and weak phylogeographic structure in domestic goats. Proceedings of the National Academy of Sciences. 2001;98: 5927–5932. doi:10.1073/pnas.091591198

13. Naderi S, Rezaei H-R, Taberlet P, Zundel S, Rafat S-A, Naghash H-R, et al. Large-Scale Mitochondrial DNA Analysis of the Domestic Goat Reveals Six Haplogroups with High Diversity. PLOS ONE. 2007;2: e1012. doi:10.1371/journal.pone.0001012

14. Naderi S, Rezaei H-R, Pompanon F, Blum MGB, Negrini R, Naghash H-R, et al. The goat domestication process inferred from large-scale mitochondrial DNA analysis of wild and domestic individuals. Proceedings of the National Academy of Sciences. 2008;105: 17659– 17664. doi:10.1073/pnas.0804782105

15. Zhao Y, Zhao R, Zhao Z, Xu H, Zhao E, Zhang J. Genetic diversity and molecular phylogeography of Chinese domestic goats by large-scale mitochondrial DNA analysis. Mol Biol Rep. 2014;41: 3695–3704. doi:10.1007/s11033-014-3234-2

16. Peng W, Zhang Y, Gao L, Feng C, Yang Y, Li B, et al. Analysis of World-Scale Mitochondrial DNA Reveals the Origin and Migration Route of East Asia Goats. Front Genet. 2022;13. doi:10.3389/fgene.2022.796979

17. VarGoats Consortium, Nijman IJ, Rosen BD, Bardou P, Faraut T, Cumer T, et al. Geographical contrasts of Y-chromosomal haplogroups from wild and domestic goats reveal ancient migrations and recent introgressions. Molecular Ecology. 2022;31: 4364– 4380. doi:10.1111/mec.16579

18. Pereira F, Carneiro J, Soares P, Maciel S, Nejmeddine F, Lenstra JA, et al. A multiplex primer extension assay for the rapid identification of paternal lineages in domestic goat (Capra hircus): Laying the foundations for a detailed caprine Y chromosome phylogeny. Molecular Phylogenetics and Evolution. 2008;49: 663–668. doi:10.1016/j.ympev.2008.08.026

19. Pereira F, Queiros S, Gusmao L, Nijman IJ, Cuppen E, Lenstra JA, et al. Tracing the History of Goat Pastoralism: New Clues from Mitochondrial and Y Chromosome DNA in North Africa. Molecular Biology and Evolution. 2009;26: 2765–2773. doi:10.1093/molbev/msp200

20. Tosser-Klopp G, Bardou P, Bouchez O, Cabau C, Crooijmans R, Dong Y, et al. Design and Characterization of a 52K SNP Chip for Goats. Liu Z, editor. PLoS ONE. 2014;9: e86227. doi:10.1371/journal.pone.0086227

21. Berihulay H, Li Y, Liu X, Gebreselassie G, Islam R, Liu W, et al. Genetic diversity and population structure in multiple Chinese goat populations using a SNP panel. Animal Genetics. 2019;50: 242–249. doi:10.1111/age.12776

22. Deniskova TE, Dotsev AV, Selionova MI, Reyer H, Sölkner J, Fornara MS, et al. SNP-Based Genotyping Provides Insight Into the West Asian Origin of Russian Local Goats. Front Genet. 2021;12. doi:10.3389/fgene.2021.708740

23. Kumar C, Song S, Dewani P, Kumar M, Parkash O, Ma Y, et al. Population structure, genetic diversity and selection signatures within seven indigenous Pakistani goat populations. Animal Genetics. 2018;49: 592–604. doi:10.1111/age.12722

24. Le SV, de las Heras-Saldana S, Alexandri P, Olmo L, Walkden-Brown SW, van der Werf JHJ. Genetic diversity, population structure and origin of the native goats in Central Laos. Journal of Animal Breeding and Genetics. 2024;141: 531–549. doi:10.1111/jbg.12862

25. Senczuk G, Macrì M, Di Civita M, Mastrangelo S, Del Rosario Fresno M, Capote J, et al. The demographic history and adaptation of Canarian goat breeds to environmental conditions through the use of genome-wide SNP data. Genet Sel Evol. 2024;56: 2. doi:10.1186/s12711-023-00869-0

26. Yonezawa T, Wu J, Masuko R, Iso K, Nomura N, Tabata R, et al. Genome-wide variation reveal that goats were introduced into Asia via multiple migrations. Research Square [Preprint]. 2025 [cited 2025 June 4]. doi: 10.21203/rs.3.rs-6559993/v1

27. Denoyelle L, Talouarn E, Bardou P, Colli L, Alberti A, Danchin C, et al. VarGoats project: a dataset of 1159 whole-genome sequences to dissect Capra hircus global diversity. Genetics Selection Evolution. 2021;53: 86. doi:10.1186/s12711-021-00659-6

28. Mbeki L, van Rossum M. Private slave trade in the Dutch Indian Ocean world: a study into the networks and backgrounds of the slavers and the enslaved in South Asia and South Africa. Slavery & Abolition. 2017;38: 95–116. doi:10.1080/0144039X.2016.1159004

29. Ricklefs MC, Lockhart B, Lau A. A New History of Southeast Asia. Bloomsbury Academic; 2010.

30. Skowronek RK. The Spanish Philippines: Archaeological Perspectives on Colonial Economics and Society. International Journal of Historical Archaeology. 1998;2: 45–71. doi:10.1023/A:1022662213844

31. Mannen H, Iso K, Kawaguchi F, Sasazaki S, Yonezawa T, Dagong MIA, et al. Indonesian native goats ( *Capra hircus*) reveal highest genetic frequency of mitochondrial DNA haplogroup B in the world. Anim Sci J. 2020;91. doi:10.1111/asj.13485

32. Masuko R, Ayin, Masaoka M, Kawaguchi F, Sasazaki S, Dagong MIA, et al. Maternal and paternal lineage analysis of Island Southeast Asian goats reveals continental propagation routes and introgression through the Indian ocean. Sci Rep. 2025;15: 9411. doi:10.1038/s41598-025-93651-9

33. Chokoe TC, Hadebe K, Muchadeyi FC, Nephawe KA, Dzomba EF, Mphahlele TD, et al. Conservation status and historical relatedness of South African communal indigenous goat populations using a genome-wide single-nucleotide polymorphism marker. Frontiers in Genetics. 2022;13. Available: https://www.frontiersin.org/articles/10.3389/fgene.2022.909472

34. Purcell S, Neale B, Todd-Brown K, Thomas L, Ferreira MAR, Bender D, et al. PLINK: A Tool Set for Whole-Genome Association and Population-Based Linkage Analyses. The American Journal of Human Genetics. 2007;81: 559–575. doi:10.1086/519795

35. Daly KG, Mattiangeli V, Hare AJ, Davoudi H, Fathi H, Doost SB, et al. Herded and hunted goat genomes from the dawn of domestication in the Zagros Mountains. Proceedings of the National Academy of Sciences. 2021;118: e2100901118. doi:10.1073/pnas.2100901118

36. Peters J, von den Driesch A, Helmer D, Saña Segui M. Early Animal Husbandry in the Northern Levant. Paléorient. 1999;25: 27–48.

37. Zeder MA, Hesse B. The Initial Domestication of Goats (*Capra hircus*) in the Zagros Mountains 10,000 Years Ago. Science. 2000;287: 2254–2257. doi:10.1126/science.287.5461.2254

38. R Core Team. R: A Language and Environment for Statistical Computing. Vienna, Austria: R Foundation for Statistical Computing; 2023. Available: https://www.R-project.org/

39. Alexander DH, Novembre J, Lange K. Fast model-based estimation of ancestry in unrelated individuals. Genome Res. 2009;19: 1655–1664. doi:10.1101/gr.094052.109

40. Huson DH, Bryant D. The SplitsTree App: interactive analysis and visualization using phylogenetic trees and networks. Nat Methods. 2024;21: 1773–1774. doi:10.1038/s41592-024-02406-3

41. Pickrell JK, Pritchard JK. Inference of Population Splits and Mixtures from Genome-Wide Allele Frequency Data. PLOS Genetics. 2012;8: e1002967. doi:10.1371/journal.pgen.1002967

42. Maier R, Flegontov P, Flegontova O, Işıldak U, Changmai P, Reich D. On the limits of fitting complex models of population history to f-statistics. Nordborg M, Przeworski M, Balding D, Wiuf C, editors. eLife. 2023;12: e85492. doi:10.7554/eLife.85492

43. Patterson N, Moorjani P, Luo Y, Mallick S, Rohland N, Zhan Y, et al. Ancient Admixture in Human History. Genetics. 2012;192: 1065–1093. doi:10.1534/genetics.112.145037

44. Porter V, Alderson L, Hall SJ, Sponenberg DP. Mason’s world encyclopedia of livestock breeds and breeding, 2 Volume Pack. Cabi; 2016.

45. Lewald KM, Abrieux A, Wilson DA, Lee Y, Conner WR, Andreazza F, et al. Population genomics of Drosophila suzukii reveal longitudinal population structure and signals of migrations in and out of the continental United States. G3 Genes|Genomes|Genetics. 2021;11: jkab343. doi:10.1093/g3journal/jkab343

46. Petretto E, Dettori ML, Luigi-Sierra MG, Noce A, Pazzola M, Vacca GM, et al. Investigating the footprint of post-domestication dispersal on the diversity of modern European, African and Asian goats. Genet Sel Evol. 2024;56: 55. doi:10.1186/s12711-024-00923-5

47. Lv F-H, Cao Y-H, Liu G-J, Luo L-Y, Lu R, Liu M-J, et al. Whole-Genome Resequencing of Worldwide Wild and Domestic Sheep Elucidates Genetic Diversity, Introgression, and Agronomically Important Loci. Molecular Biology and Evolution. 2022;39: msab353. doi:10.1093/molbev/msab353

48. Manunza A, Ramirez-Diaz J, Cozzi P, Lazzari B, Tosser-Klopp G, Servin B, et al. Genetic diversity and historical demography of underutilised goat breeds in North-Western Europe. Sci Rep. 2023;13: 20728. doi:10.1038/s41598-023-48005-8

49. Mwacharo JM, Bjørnstad G, Mobegi V, Nomura K, Hanada H, Amano T, et al. Mitochondrial DNA reveals multiple introductions of domestic chicken in East Africa. Molecular Phylogenetics and Evolution. 2011;58: 374–382. doi:10.1016/j.ympev.2010.11.027

50. Casimiro TM, Borges MO. Life on Board Portuguese Ships in the 16th–18th Centuries: Theorizing Households through History and Archaeology. Heritage. 2023;6: 2020–2037. doi:10.3390/heritage6020109

51. Lobo J, Costa MG da, Beckingham CF (Charles F, Lockhart DM. Itinerário, e outros escritos inéditos. Livraria Civilização; 1971. Available: https://cir.nii.ac.jp/crid/1130282270486239616

52. Chias P, Abad T. Colonial Urban Planning and Land Structures in the Philippines, 1521-1898. Journal of Asian Architecture and Building Engineering. 2012;11: 9–16. doi:10.3130/jaabe.11.9

53. Zuhdi S. Shipping Routes and Spice Trade in Southeast Sulawesi in the 17th and 18th Century. Journal of Maritime Studies and National Integration. 2018;2: 31–44. doi:10.14710/jmsni.v2i1.3100

54. Fourie J, Jansen A, Siebrits K. Public finances under private company rule: The Dutch Cape Colony (1652–1795). New Contree. 2013;68: 21. doi:10.4102/nc.v68i0.277

55. Nguluma A, Kyallo M, Tarekegn GM, Loina R, Nziku Z, Chenyambuga S, et al. Mitochondrial DNA D-loop sequence analysis reveals high variation and multiple maternal origins of indigenous Tanzanian goat populations. Ecol Evol. 2021;11: 15961–15971. doi:10.1002/ece3.8265

